# Revised diffusion law permits quantitative nanoscale characterization of membrane organization

**DOI:** 10.1101/2024.11.01.621464

**Authors:** Barbora Svobodová, David Šťastný, Hans Blom, Ilya Mikhalyov, Natalia Gretskaya, Alena Balleková, Erdinc Sezgin, Martin Hof, Radek Šachl

**Author notes:** **Correspondence:** Radek Šachl, Martin Hof, Erdinc Sezgin.

## Abstract

Formation of functional nanoscopic domains is an inherent property of plasma membranes. Stimulated emission depletion combined with fluorescence correlation spectroscopy (STED-FCS) has been used to identify such domains, however, the information obtained by STED-FCS has been limited to presence of such domains while crucial parameters have not been accessible, such as size (*R*_d_), the fraction of occupied membrane surface (*f*), in-membrane lipid diffusion inside (*D*_in_) and outside (*D*_out_) the nanodomains as well as their self-diffusion (*D*_d_). Here, based on a revision of the ‘diffusion law’, we present an approach to retrieve these five parameters from STED-FCS data. We verify that approach on ganglioside nanodomains in giant unilamellar vesicles (GUVs), validating the Saffman-Delbrück assumption for *D*_d_. We examined STED-FCS data in both plasma membranes of living PtK2 cells and in giant plasma membrane vesicles (GPMVs) and present a quantitative framework for molecular diffusion modes in biological membranes.

## Manuscript

Thanks to its excellent spatiotemporal resolution, STED-FCS allows adequate diffusion law plot analysis [1] at the nanoscale for which it has become a promising tool to study lipid dynamics within nanoscopically heterogeneous membranes [2]–[13]. In this approach, the apparent diffusion coefficient (*D*) of a fluorescently labelled molecule diffusing in the membrane is typically plotted against the spot radius (*w*) (**Figure 1 A,B)** [14], [15]. For optimal sensitivity, the spot radius should approach or be smaller than the size of the obstacles being studied. In case of nanosized membrane obstacles, this condition leads to what we call STED-FCS diffusion law plot dependencies, where the diffusion coefficient is measured for waists ranging between 20-250 nm. These diffusion law plots typically show a specific pattern, depending on the type of interactions and obstacles in the lipid bilayer (**Figure 1A)**. The complexity of lipid dynamics generates exceptions to this rule, though, and occasionally one can get the same pattern for several diffusion modes [16].

**Figure1:**
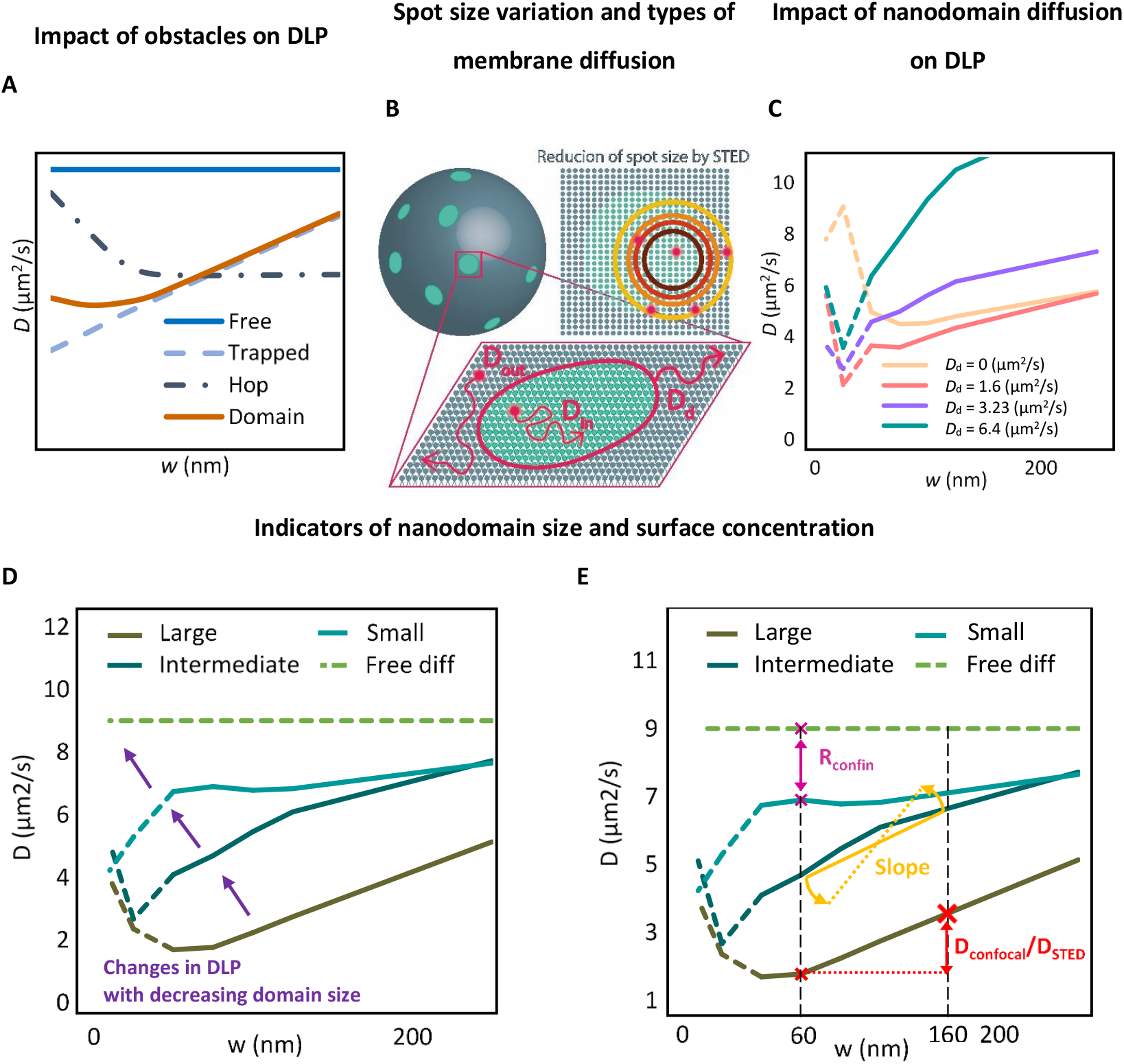
(A) Diffusion law plots (DLP) depicting the relationship between the probe diffusion coefficient (*D*) and the radius of the illuminated focal spot (*w*). The shape of these plots reflects the type of obstacles in the membranes: Free diffusion in homogeneous membrane areas shows a constant *D*, independent of the spot radius (solid blue line); Trapped diffusion occurs when molecules are temporarily immobilized, resulting in a decreasing *D* as *w* decreases (dashed light blue line); Hop diffusion involves movement within a meshwork structure, with fast diffusion over short distances and impeded diffusion over longer ones (dashed dotted dark blue line). The diffusion in the presence of immobile nanodomains leads to a gradual decrease of *D* with decreasing *w* and flattening out of this dependence for small waist radii (solid orange line). (B) In STED-FCS, the effective spot size is reduced by applying stronger STED laser pulses. For membranes with mobile nanodomains, three diffusion types can exist: probe diffusion inside (*D*_in_) and outside (*D*_out_) nanodomains, and diffusion of nanodomains themselves (*D*_d_). (C) Diffusion law plots for nanodomains with increasing mobility: static nanodomains (*D*_d_ = 0) and mobile nanodomains (*D*_d_ = 1.6 – 6.4 µm^2^/s). (D,E) Indicators of nanodomain properties from STED-FCS data and diffusion law plots: (D) Shape of diffusion law plots, sensitive to nanodomain presence and their physical-chemical features. (E) Characteristic ratiometric parameters involving: *D*_confocal_/*D*_STED_ diffusion coefficient ratio determined for two key waists *w*_confocal_ = 160 and *w*_STED_ = 60 nm (depicted in red); confinement ratio *R*_confin_ = *D*(no domains)/*D*_60_ expressing the reduction in diffusion rate attributed to nanodomain formation (depicted in magenta) and the slope 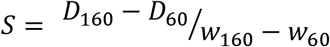 calculated for the waist range 60 – 160 nm (depicted in yellow). For a homogeneous membrane, *D*_160_/*D*_60_ = *R*_confin_ = *S* = 1. With mobile nanodomains, these values are greater than 1.

Interpretations of diffusion data have so far relied on assumptions that are not always universally accepted. Particularly, in cellular membranes with nanoscopic domains, these domains were assumed to be static despite their generally dynamic nature, which may affect the interpretation of the results (**Figure 1C**)[1], [16]–[19]. In this work, we incorporate nanodomain mobility into the analysis of STED-FCS diffusion law plots. The analysis of *in-silico* generated STED-FCS diffusion law plots reveals characteristic shapes with distinct fingerprints that indicate the size (*R*_d_) and surface concentration (*f*) of nanodomains. Next, we verify this analysis on STED-FCS data collected on a well-defined model system comprising membranes containing ganglioside GM_1_ nanodomains of known *R*_d_ and *f*. In the final step, we revisit previously published diffusion law plots for fluorescently labelled ganglioside GM_1_ in both cellular plasma membranes and giant plasma membrane vesicles [18]. This reassessment not only confirms the validity of our experimental approach but also uncovers previously inaccessible information in STED-FCS diffusion law plots, significantly enhancing the interpretative framework for STED-FCS data in the context of mobile membrane nanodomains.

To exploit the full potential of STED-FCS diffusion law plots and ensure their unbiased interpretation, our primary goal was to pinpoint distinctive trends within these plots that could signify key nanodomain features such as *R*_d_ and *f*. We thus initiated our study by generating a series of these dependencies *in silico*. This involved exploring various combinations of parameter values that describe the diffusion of fluorescent lipid probes in the presence of mobile nanodomains (see **Supporting Information (SI)** and **Figure 3**). Specifically, we kept the diffusion coefficient of lipids outside the nanodomains (*D*_out_) using previously determined values for labelled GM_1_ in GUVs.[20] The sizes of the nanodomains (*R*_d_) were chosen to match the sizes commonly encountered in biological membranes: 1) small nanodomains with a radius of 25 nm; 2) intermediate-sized nanodomains with a radius of 75 nm; and large nanodomains with a radius of 120 nm. The surface concentration of nanodomains (*f*), was chosen to range from 10 to 50%, with the upper limit approximately determined by the maximum number of nanodomains that can still be placed side by side so as not to overlap. Regarding 2-dimensional nanodomain movement, our simulations exclusively considered nanodomains moving with a diffusion coefficient (*D*_d_) calculated using the Saffman-Delbrück model[21], a choice supported by our later experimental findings in this work. Additionally, we anticipated that the shape of the diffusion law plots could be significantly influenced by the partition coefficient of probes in the nanodomains (*K*_d_), as well as the probe diffusion rate within nanodomains (*D*_in_). Consequently, we expanded the set of generated diffusion law plots to include these dependencies as well. The results obtained are summarized in **Figure 2**, with a comprehensive set of generated diffusion law plot dependencies available in the extended **Figure S1** in the **Supporting Information** file.

**Figure 2:**
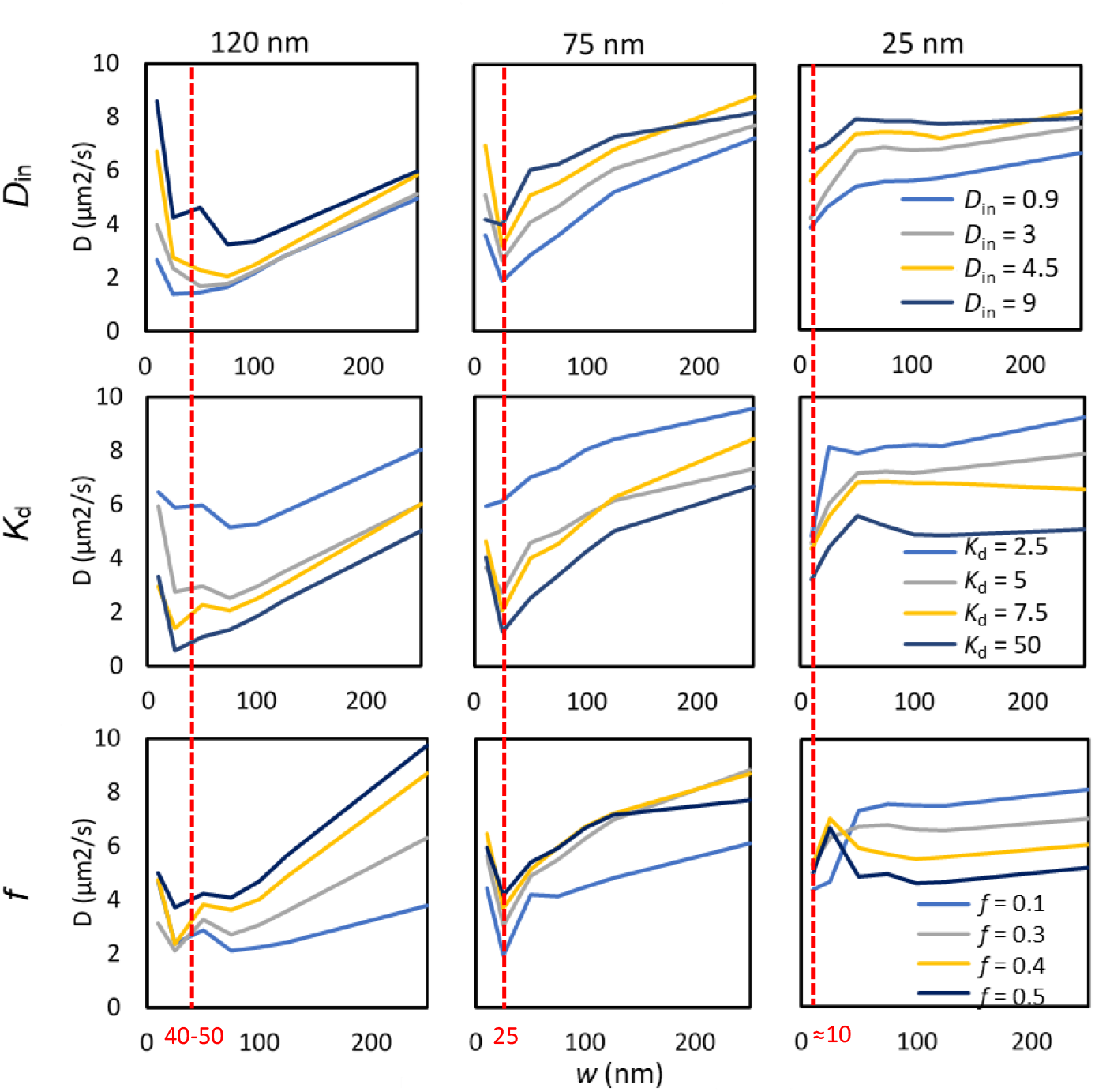
Computationally generated STED-FCS diffusion law plots illustrating mobile nanodomains with *D*_d_ modelled according to the Saffman-Delbrück model. Plots are shown for three nanodomain sizes: large (left column), intermediate (middle column) and small (right column). The impact of the probe diffusion coefficient within the nanodomains (*D*_in_) is depicted in the upper row, the probe distribution coefficient between the nanodomains and the surroundings (*K*_d_) in the middle row, and the area fraction (*f*) occupied by nanodomains in the lower row. If not stated otherwise, *D*_in_ = 4.5 µm^2^/s, *K*_d_ =5, *f* = 0.25 *D*_out_ = 9 µm^2^/s. The red line indicates the minimum in the diffusion law plot corresponding to approximately one-third of *R*_d_. Extended data set is shown in **Figure SI1**.

**Figure 3:**
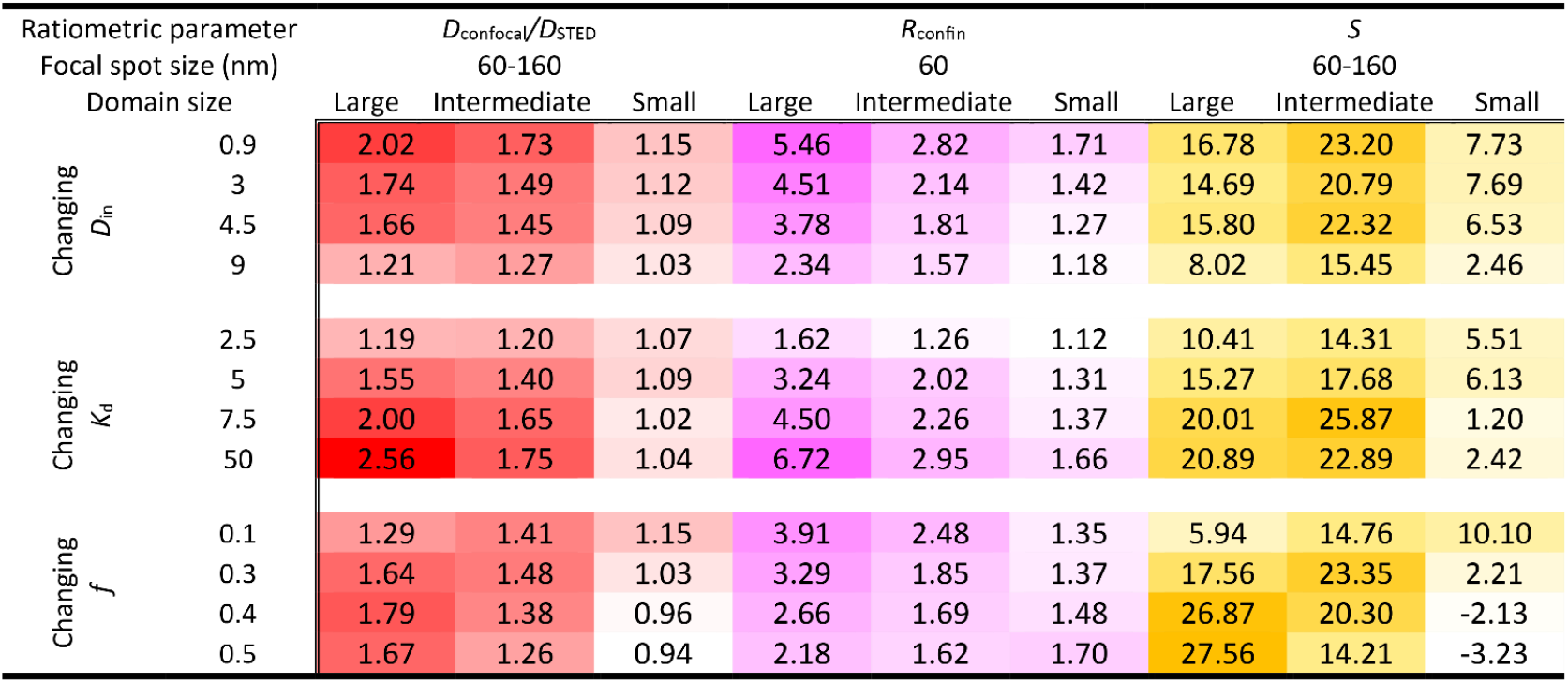
A color map of ratiometric parameters *D*_confocal_/*D*_STED_ (red), *R*_confin_ (violet), and *S* (yellow) used as indicators of nanodomain size and fraction. The parameters were calculated for large (*R*_d_ = 120 nm), intermediate (*R*_d_ = 75 nm), and small nanodomains (*R*_d_ = 25 nm), based on the probe diffusion coefficient within the nanodomains (*D*_in_), probe partition coefficient between nanodomains and surroundings (*K*_d_), and fraction of nanodomains in the membrane (*f*). The colour code illustrates a gradual change from low (light) to high (dark) values. If not stated otherwise, *D*_in_ = 4.5 µm^2^/s, *D*_out_ = 9 µm^2^/s, *K*_d_ = 5 and *f* = 0.25. An extended version of this figure is presented in **Figure S1**.

Analysing trends in these dependencies revealed several indicators of nanodomain properties (**Figure 1D,E**). The diffusion law plots exhibit a characteristic shape reminiscent of an asymmetric funnel profile, with a broader shoulder extending towards larger focal waists (**Figure 2**). The minimum point of this funnel appears to correspond to approximately one-third of the nanodomain radius *R*_d_ (highlighted by red dashed lines in **Figure 2**), while its depth is determined by the lipid probes’ affinity for the nanodomains (*K*_d_) and their diffusion rate within them (*D*_in_). Stronger entrapment and retardation of probe movement within nanodomains lead to deeper minima in the plots. Meanwhile, *R*_d_ dictates the extent of this characteristic shape captured our in-silico analysis. Whereas the large nanodomains (*R*_d_ = 120 nm) exhibit a strong funnel profile dependence that is missing the final plateau in the waist range 20-250 nm, the small ones (*R*_d_ = 25 nm) only reveal the final plateau of the extended funnel arm. This analysis thus highlights characteristic fingerprinting of nanodomain size in the diffusion law plots, mirrored in the position of the minimum in the funnel-shaped dependence and the segment of this dependence captured (**Figure 4A**). In this context, it is also worth noting a similar funnel-shaped profile trend in the diffusion law plots linking the anomalous factor *α*, determined by fitting individual STED-FCS autocorrelation functions by a model accounting for anomalous diffusion **(Eq. 2** in **Supporting Information**), and the size of the confocal spot *w* (**Figure S2**).

**Figure 4:**
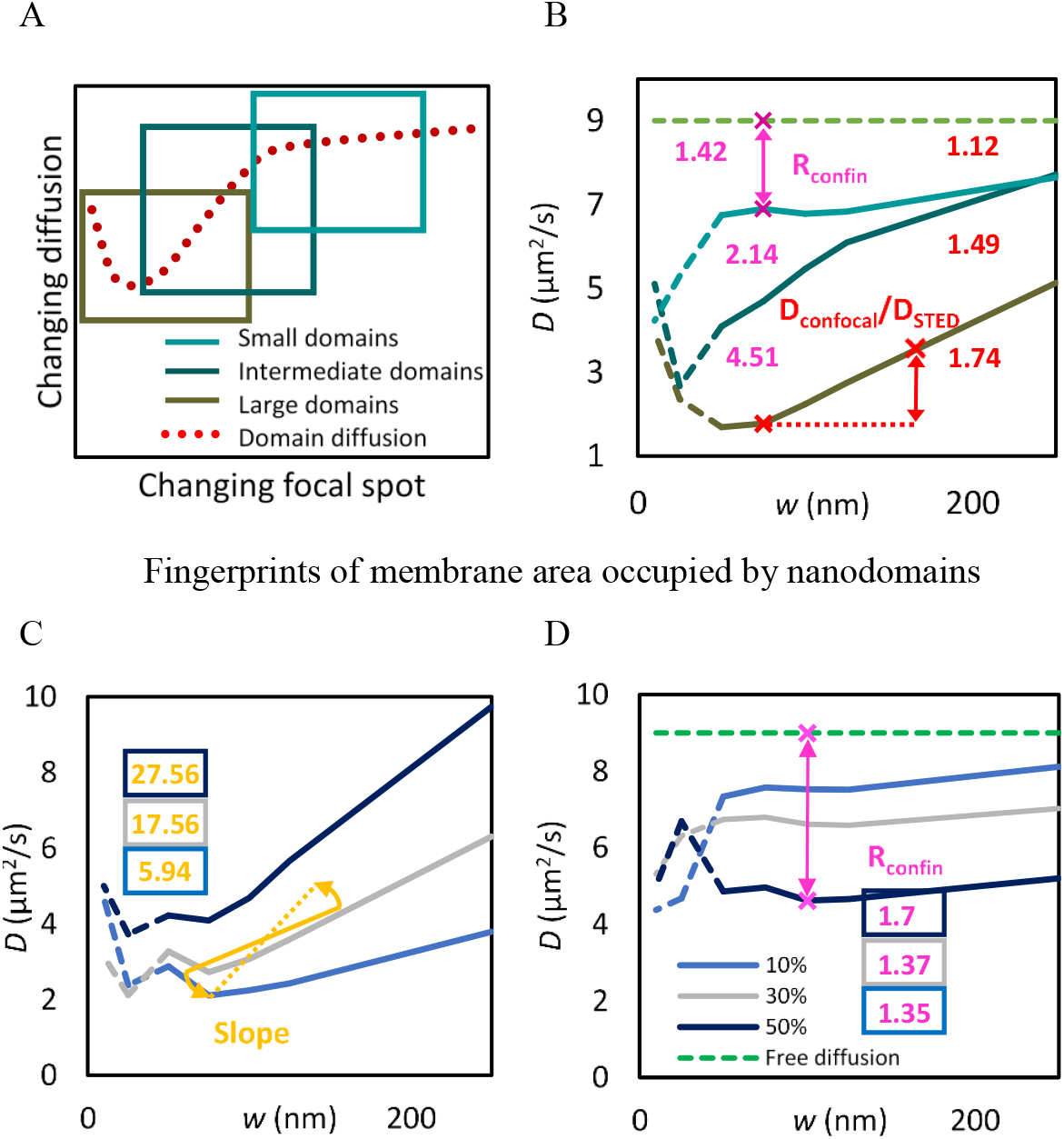
(AB) Identified fingerprints indicative of nanodomain size. (A) The diffusion law plot in the presence of moving nanodomains exhibits a characteristic shape resembling an asymmetric funnel. The nanodomain size determines the extent of this characteristic shape observed in the experiment. Large nanodomains (*R*_d_ = 120 nm) display a funnel-like dependence lacking a final plateau in the waist range of 20-250 nm (dark brown), intermediate-sized nanodomains (*R*_d_ = 75 nm) show a rising portion of the funnel that gradually levels off (dark green), while small nanodomains ((*R*_d_ = 25 nm) reveal only the final plateau of the extended funnel shape (cyan). (B) Ratiometric indicators of nanodomain size: *D*_confocal_/*D*_STED_ ratio (depicted in red) derived from the probe diffusion coefficients *D* determined for *w* = 160 nm and *w* = 60 nm. *D*_confocal_/*D*_STED_ > 1.8 indicates the presence of large domains, *D*_confocal_/*D*_STED_ ∈ (1.2; 1.6) suggests intermediate-sized nanodomains, while *D*_confocal_/*D*_STED_ < 1.2 suggests small nanodomains or a homogeneous membrane. Confinement ratio *R*_confin_ = *D*_(no entrap.)_/*D*_60_ (depicted in magenta), where *D*(no entrap.) is the diffusion coefficient of a probe showing no entrapment in nanodomains, and *D*_60_ is the diffusion coefficient of a probe confined within nanodomains and measured with the beam spot radius *w* = 60 nm. *R*_confin_ > 2.8 indicates the presence of large domains, while *R*_confin_ ∈ (1.7; 2.8) suggests intermediate-sized nanodomains. *R*_confin_ < 1.7 suggests small nanodomains or a homogeneous membrane. (C,D) Identified fingerprints indicative of nanodomain fraction *f* shown for both large (C) and small nanodomains (D). *K*_d_ = 5 and *D*_in_ = 4.5 (C) In the case of large nanodomains, the slope of the diffusion law plot dependence calculated as 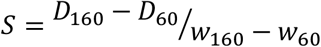 emerges as the primary indicator of *f*. This slope exhibits a low value when *f* = 0.1 (*S* ≈ 5), increasing to as high as *S* ≈ 27 for *f* = 0.5. (D) In the case of small nanodomains, *f* is primarily indicated by *R*_confin_ increasing from *R*_confin_ ≈ 1.3 for *f* = 0.1 to *R*_confin_ ≈ 1.7 for *f* = 0.5. As anticipated, higher *f* results in more pronounced confinement.

In order to simplify the analysis of STED-FCS data, Sezgin et al introduced a simplified approach centred on the analysis of a single ratiometric parameter *D*_confocal_/*D*_STED_ [12]. This diffusion coefficient ratio is derived from the probe diffusion coefficients *D* determined for two focal spot sizes: *w* = 160 nm, denoted as *D*_confocal_ (representing the confocal spot size in our case), and *w* = 60 nm, denoted as *D*_STED_ (a waist size typically achievable with STED microscopy) (see also **Figure 1E** for the definition of this parameter). Widely adopted in literature, the parameter *D*_confocal_/*D*_STED_ acts as a tentative indicator of nanoscale membrane heterogeneity, essentially reflecting the rate of diffusion retardation relative to the focal spot size *w* [3], [12], [15], [18], [22]. This ratio is consolidated in a part of **Figure 3** and **Figure 3B**, supplementing the diffusion dependencies illustrated in **Figure 2**. In a homogeneous membrane devoid of obstacles *D*_confocal_/*D*_STED_ = 1.

Notably, **Figure 3** demonstrates that *D*_confocal_/*D*_STED_ also holds strong predictive value regarding the nanodomain size. Specifically, an *D*_confocal_/*D*_STED_value exceeding 1.8 robustly indicates the presence of large domains, except in rare instances of exceptionally high *K*_d_ or extremely slow diffusion within the nanodomains. Conversely, an *D*_confocal_/*D*_STED_ value falling within the range of 1.2-1.6 suggests intermediate-sized nanodomains, while an *D*_confocal_/*D*_STED_ value below 1.2 suggests small nanodomains or a homogeneous membrane without obvious barriers.

To differentiate between the latter cases, one can assess the reduction in diffusion rate attributed to nanodomain formation using an additional parameter defined as: *R*_confin_ = *D*_no entrap_/*D*_60_ and called the confinement ratio (**Figure 4B**) (see also **Figure 1E** for the definition of this parameter). Even with minimal probe entrapment in nanodomains, this ratio notably deviates from 1, even in the presence of small domains (**Figure 3** and more detailed **Table S1**). Although it appears to offer superior sensitivity to nanodomain size as compared to *D*_confocal_/*D*_STED_, obtaining it necessitates an additional measurement of *D*_no entrap_ in a membrane environment where no probe entrapment occurs. This can be done either in membranes devoid of obstacles or using fluorescent probes that do not become trapped in nanodomains. Overall, our in-silico analysis shows that STED-FCS diffusion law plots generated for mobile nanodomains contain a variety of signatures that are indicative of the presence and approximate size of the detected nanodomains (**Figure 4**).

In fact, our in-silico analysis also predicts the shape of the diffusion law plot dependencies to be largely influenced by the surface concentration of nanodomains *f*. However, interpreting these dependencies becomes more complex because the response of the diffusion law plot to *f* relies heavily on *R*_d_, while the influence of either *K*_d_ or *D*_in_ is relatively minor (**Figure 2**). Fundamentally, two distinct scenarios emerge in diffusion law plots, each offering unique fingerprints of the nanodomain fraction, contingent upon nanodomain size: In the case of large nanodomains, *f* is distinctly determined by the slope (*S*) of the respective dependence 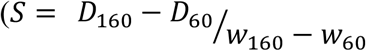, (see **Figure 1E** for the definition of this parameter and **Figure 4C** for the revealed trends). Conversely, for small nanodomains, the confinement ratio emerges as the primary indicator of *f* (**Figure 4D**).

To experimentally validate and utilize the nanodomain indicators identified through computational simulations, we turned in the next step to an inherently nanoscopically heterogeneous system of GUVs containing physiologically relevant amounts of ganglioside GM_1_[23]–[30]. We recorded STED-FCS diffusion law plots using GM_1_ labelled in the headgroup with Atto565 as a lipid tracer (referred to as GM_1_-Atto565). Gangliosides exhibit a pronounced tendency to segregate into lipid nanodomains spanning 10-120 nm in radius, influenced by the specific ganglioside type and the composition of the surrounding environment [23]– [26], [31], [32]. Recently, we also characterized this system in detail using the Monte-Carlo Förster Resonance Energy Transfer (MC-FRET) technique, which determines both the radius and surface concentration of these formed nanodomains [33]–[36]. Consequently, we possessed a well-defined model system with ganglioside nanodomains of known *R*_d_ and *f* [24], [25]. Our primary objective extended beyond merely applying identified qualitative indicators to an experimental system featuring mobile nanodomains. We sought to conduct a thorough quantitative analysis of STED-FCS diffusion law plots, a task not previously undertaken for such systems.

For our experiments, we chose GUVs containing two different types of nanodomains: a) large nanodomains with *R*_d_ = 120 nm and nanodomain surface coverage *f* = 0.5 [24], [25] formed in 1,2-dioleoyl-*sn*-glycero-3-phosphocholine (DOPC)/cholesterol (Chol)/GM_1_ bovine brain sodium salt (GM_1_) (75/25/4 mol%) mixtures, where the nanodomain size matched perfectly the size of large nanodomains characterized *in-silico*; and b) small nanodomains formed in DOPC/N-stearoyl-D-erythro-sphingosylphosphorylcholine (SM)/GM_1_ (90/10/4 mol%) mixtures that according to the previously performed MC-FRET analysis had a radius of 23 nm and occupied 47 % [24], [25] of the bilayer surface. As a result, these nanodomains had a size comparable to smallest waist diameter we could accomplish in our experimental setup 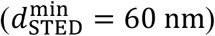 resembled in size small nanodomains used in the simulations.

The experimental results in **Figure 5A** display STED-FCS diffusion law plots for both large and small GM_1_ nanodomains utilizing GM_1_ labelled in the headgroup with Atto565 as a lipid tracer (referred to as GM_1_-Atto565). With this fluorescent label, we achieved STED focal spot sizes ranging from 60 to 160 nm. This spatial range allowed for a comprehensive quantitative analysis, although it was not large enough to fully capture the funnel-shaped dependency, even for large nanodomains (compare **Figure 4A** with **Figure 5A**).

**Figure 5:**
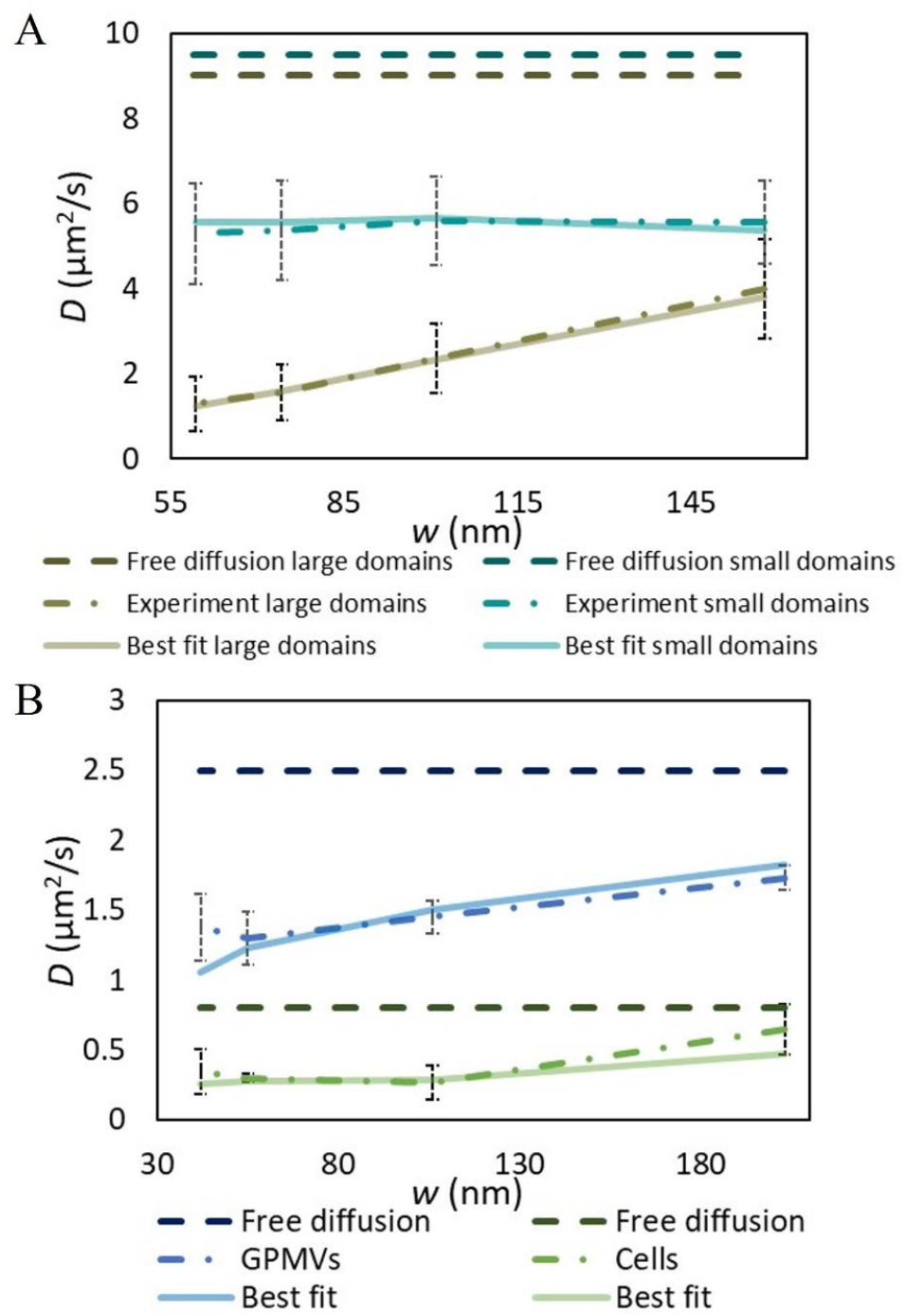
Experimentally obtained STED-FCS diffusion dependencies (dashed-dotted lines) and their best fits for GM_1_-Atto565 diffusion in: (A) DOPC/Chol/GM_1_ (75/25/4) GUVs containing large nanodomains (*R*_d_ = 120 nm) and DOPC/SM/GM_1_ (90/10/4) GUVs containing small nanodomains (*R*_d_ = 23 nm). (B) Experimentally measured diffusion dependencies (adopted from [18]) and their best fits for GM_1_-Atto647N diffusing: in the plasma membranes of PtK2 cells (green curves) and in GPMVs made from these cells (blue curves). In this case, the quantitative analysis of STED-FCS diffusion law plots was used to determine *R*_d_, *f*, and *D*_in_. For comparison, limiting cases of free diffusion with a waist-independent diffusion coefficient value corresponding to *D*_out_ (solid green) are shown in all panels.

Importantly, the diffusion law plots for both large and small nanodomains exhibited characteristic features consistent with our simulations of probe diffusion in the presence of mobile nanodomains. Specifically, the diffusion law plot pattern for large nanodomains showed a sharp increase without fully reaching a plateau (compare **Figure 4A with Figure 5A**). Moreover, the introduction of the simplified ratiometric parameters *D*_160_/*D*_60_ = 3.07 and *R*_confin_ = 6.94 confirmed the presence of nanodomains with an estimated radius of approximately 120 nm (**Figure 4B** and **Figure 3**). Evaluating the diffusion law slope parameter, *S* = 27.35, suggested a surface concentration of nanodomains with a fraction *f* greater than 0.3 (**Figure 4C and Figure 3**), aligning with our assumptions based on MC-FRET experiments [24], [25]. In contrast, the diffusion law plot pattern for small nanodomains appeared flat, with the diffusion ratio *D*_160_/*D*_60_ = 1.04, indicating the presence of either very small nanodomains or a homogeneous membrane (**Figure 4B** with **Figure 3**). A homogeneous membrane was unlikely given that the confinement ratio *R*_confin_ = 1.79 deviated significantly from 1. This high *R*_confin_ value further suggests a high surface concentration of very small nanodomains (*f* = 0.4), consistent with our previous MC-FRET results (compare **Figure 4D** with **Figure 3**) [24]. In summary, the observed patterns in our experimentally recorded STED - FCS diffusion law plots, as depicted in **Figure 5**, closely match the fingerprints predicted by our MC simulations. This alignment validates the robust application of STED-FCS diffusion law plots to nanoscopically heterogeneous membrane systems featuring dynamic nanodomains, paving the way for quantitative analyses of diffusion law plot dependencies.

To quantitatively analyse the STED-FCS diffusion law plots in the presence of GM_1_ nanodomains (**Figure 5A**), we generated an extended set of diffusion law plot dependencies using various combinations of simulation input parameters (To access the library containing a complete set of generated diffusion law plots click on the following link: **[data repository link forthcoming]**). Our goal was to identify the combination of these parameters that best matched the experimental data, as indicated by the lowest values of the chi-squared parameter.

Our analysis specifically targeted the STED-FCS diffusion law plots for GM_1_ nanodomains within DOPC/Chol/ GM_1_ (75/25/4 mol%) or DOPC/SM/GM_1_ (90/10/4 mol%) GUVs, particularly examining the diffusion law plots for large and small nanodomains as depicted in **Figure 5**. Since the parameters *R*_d_ and *f* were determined independently using MC-FRET, and *D*_out_ was experimentally determined by FCS (9 μm^2^/s and 9.5 μm^2^/s for free diffusion in DOPC/Chol (75/25) and in DOPC/SM (90/10), respectively [20]), our optimization efforts centred on refining *D*_d_, *D*_in_, and *K*_d_. In the final stage, slight adjustments were made to the best-matching curve to optimize its vertical positioning, involving fine-tuning of the fraction *f* (**Figure 5A**). This iterative process enabled us to determine optimal values of *D*_d_, *D*_in_, and *K*_d_ for both large and small GM_1_ nanodomains. The values obtained, with *D*_d_ = 2.6 μm^2^/s for large and *D*_d_ = 4.8 μm^2^/s for small nanodomains, confirm the high mobility of GM_1_ nanodomains within GUV membranes, consistent with the Saffman-Delbrück model (implying *D*_d_ is a function of *R*_d_). Furthermore, a *K*_d_ value of 10 for both large and small nanodomains indicates that individual ganglioside molecules are predominantly localized within nanodomains, with only minimal occurrence outside these domains in the membrane. This finding underscores GM_1_’s strong tendency to segregate spatially into nanodomains[23]–[25], [29], [31], [37]–[39].

Our approach also facilitates the examination of lipid diffusion rates within nanodomains, a parameter that is typically challenging to determine directly. Our analysis revealed that *D*_in_(GM_1_ − Atto565) = 4.5 μm^2^/s in both small and large nanodomains, which is only twice as slow as the diffusion coefficient recorded for homogeneous DOPC/Chol/GM_1_ (75/25/4) membranes. This indicates that the retarding effect of the packed ganglioside environment is only mild supporting the fluid and disordered character of GM_1_ nanodomains[9].

In the final comparison analysis, we applied the developed quantitative procedure to re-examine previously published STED-FCS diffusion law plots for Atto647N-labeled ganglioside GM_1_ (GM_1_-Atto647N) in both plasma membranes of living cells and in giant plasma membrane vesicles (GPMVs) lacking cytoskeleton[18]. These dependencies (**Figure 5B**) were previously used primarily as evidence for gangliosides being localized into lipid nanodomains in both cellular plasma membranes and in GPMVs, with qualitative characterization of molecule diffusion based on diffusion law plots generated for static nanodomains. Given that the diffusion of GM_1_-Atto647N, unlike sphingomyelin or other lipids, is largely insensitive to the naturally occurring cytoskeleton in cellular plasma membranes causing detected hop diffusion [18], [40], [41], this system thus appears ideal for applying our developed quantitative analysis.

Following our previously outlined STED-FCS diffusion law plot analysis procedure, we first determined the ratiometric parameters *D*_confocal_/*D*_STED_ and *R*_confin_ for both cell membranes (*D*_confocal_/*D*_STED_ = 1.38 and *R*_confin_ = 2.85) and GPMVs (*D*_160_/*D*_60_ = 1.36 and *R*_confin_ = 2.03). The analysis of these parameters with help of **Figure 4** and **Figure 3** allowed us to estimate the approximate size of the nanodomains in which GM_1_-Atto647N is entrapped. Specifically, assuming an increased affinity of GM_1_-Atto565 to the nanodomains (*K*_d_ = 5-10), GM_1_ appears to be confined into intermediate to large nanodomains with *R*_d_ ≈ 75 − 120 nm in plasma membranes, whereas in GPMVs, it aggregates into intermediate-sized nanodomains with *R*_d_ ≈ 75 nm. To refine these estimates, we performed a quantitative analysis by identifying an in-silico generated diffusion law plot from the online library that best fit the experimentally obtained dependence (**Figure 5B**, for more details see **Materials and Methods**). This analysis revealed that in plasma cell membranes, GM_1_-Atto647N is segregated into nanodomains with a radius *R*_d_ = 120 nm, *f* = 0.3 and *D*_in_ = *D*_out_/2 = 0.4 µm^2^/s. These GM_1_ nanodomain characteristics resemble those for GPMVs with *R*_d_ = 75 nm, *f* = 0.2 and *D*_in_ = *D*_out_/2 = 1.25 µm^2^/s. Importantly, these nanodomain features are also consistent with independent MC-FRET experiments performed on DOPC/Chol/SM/GM_1_ (65/25/10/4) GUVs, which revealed GM_1_ nanodomains with *R*_d_ = 99 nm and *f* = 0.5 [24], [25]. Furthermore, the results appear consistent also in terms of the recovered *D*_in_ coefficients that are consistently half the value of *D*_out_, indicating the that the fluidity of the nanodomain interior is similar to that of the surrounding membrane phase.

In summary, the revision of the diffusion law yielded in a comprehensive library of STED-FCS diffusion law plots for various conditions (see **Figure 2** and **Figure S2** and the following link: **[data repository link forthcoming]**). These plots showed an asymmetric funnel-like profile influenced by *R*_d_, *D*_d_ and *f*. The mobility of lipids within the nanodomains (*D*_in_) and their affinity for the nanodomains (*K*_d_), only accentuated this characteristic dependence. Based on these results, we identified *R*_d_ and *f* as the two key parameters shaping the final dependence, allowing for the identification of simple parameters to approximately estimate these parameters (**Figure 4**). Importantly, the previously published STED-FCS diffusion law plots [3], [18], [22], characterized by a gradual decrease in the diffusion coefficient *D* with decreasing focal spot waist radius *w* and flattening for small waist radii (**Figure 1C**), represent an incomplete dependence applicable only to large nanodomains with *R*_d_ ≥ 120 nm. The neglected increase in *D* with decreasing *w* for the smallest STED spot waists or the final plateau for the largest waists (**Figure 2**) were previously unshown. Application of our extended computational approach on GUV’s membranes with GM_1_ nanodomains, previously characterized by MC-FRET, confirm the Saffman-Delbrück model for nanodomain diffusion *D*_d_ and show that *D*_in_ to be only twice as slow as *D*_out_. These findings hold also for the diffusion of GM_1_ in membranes of GPMVs and PtK2 cells. The fact that, similar to the results for model membrane systems, fluid domains of about 120 nm size occupy a large fraction of the cell membrane (*f* = 0.3) indicate that that extensive nano-structuring represent a generic phenomenon in ganglioside containing membranes. Overall, we believe that this new quantitative framework for diffusion law analysis will allow resolving the nanoscale architecture of other macromolecules in cells, enhancing the level of detail and information obtainable from STED-FCS data.

## Author contributions

BS simulations of the diffusion law plots, involvement in all measurements and data analysis; DŠ fluorescence correlation spectroscopy measurements at Heyrovský Institute; HB support on STED-FCS microscopy; IM and NG synthesis of GM1-Atto565 fluorescent probe; AB pilot STED-FCS measurements and data analysis; ES supervision of STED-FCS experiments and STED-FCS measurements; ES, MH and RŠ study conception and design; RŠ supervision of the project; BS, MH and RŠ writing of the manuscript; revision of the manuscript by all authors.

## Acknowledgements

We thank the SciLifeLab Advanced Light Microscopy facility and National Microscopy Infrastructure (VR-RFI 2016-00968) for their support on imaging. ES is supported by Swedish Research Council Starting Grant (grant no. 2020-02682). BS would like to acknowledge support from Charles University, Faculty of Mathematics and Physics via GAUK 272923. BS and MH acknowledge GAC?R Grant 19-26854X. BS, MH and RŠ acknowledge the Advanced Multiscale Materials for Key Enabling Technologies project, supported by the Ministry of Education, Youth, and Sports of the Czech Republic, Project No. CZ.02.01.01/00/22 008/0004558, co-funded by the European Union.

## References

[1] L. Wawrezinieck, H. Rigneault, D. Marguet, and P. F. Lenne, “Fluorescence correlation spectroscopy diffusion laws to probe the submicron cell membrane organization,” Biophys. J., vol. 89, no. 6, pp. 4029–4042, 2005, doi: 10.1529/biophysj.105.067959.

[2] C. Eggeling et al., “Direct observation of the nanoscale dynamics of membrane lipids in a living cell,” Nature, vol. 457, no. 7233, pp. 1159–1162, 2009, doi: 10.1038/nature07596.

[3] E. Sezgin et al., Measuring nanoscale diffusion dynamics in cellular membranes with super-resolution STED–FCS, vol. 14, no. 4. 2019. doi: 10.1038/s41596-019-0127-9.

[4] X. W. Ng, N. Bag, and T. Wohland, “Characterization of lipid and cell membrane organization by the fluorescence correlation spectroscopy diffusion law,” Chimia (Aarau)., vol. 69, no. 3, pp. 112–119, 2015, doi: 10.2533/chimia.2015.112.

[5] N. Bag, X. W. Ng, J. Sankaran, and T. Wohland, “Spatiotemporal mapping of diffusion dynamics and organization in plasma membranes,” Methods Appl. Fluoresc., vol. 4, no. 3, 2016, doi: 10.1088/2050-6120/4/3/034003.

[6] G. Vicidomini, P. Bianchini, and A. Diaspro, “STED super-resolved microscopy,” Nat. Methods, vol. 15, no. 3, pp. 173–182, 2018, doi: 10.1038/nmeth.4593.

[7] N. K. Sarangi, C. Roobala, and J. K. Basu, “Unraveling complex nanoscale lipid dynamics in simple model biomembranes: Insights from fluorescence correlation spectroscopy in super-resolution stimulated emission depletion mode,” Methods, vol. 140–141, no. 2018, pp. 198–211, 2018, doi: 10.1016/j.ymeth.2017.11.011.

[8] A. Honigmann et al., “Scanning STED-FCS reveals spatiotemporal heterogeneity of lipid interaction in the plasma membrane of living cells,” Nat. Commun., vol. 5, no. 1, p. 5412, Nov. 2014, doi:10.1038/ncomms6412.

[9] R. Šachl et al., “On multivalent receptor activity of GM1 in cholesterol containing membranes,” Biochim. Biophys. Acta - Mol. Cell Res., vol. 1853, no. 4, pp. 850–857, 2015, doi: 10.1016/j.bbamcr.2014.07.016.

[10] F. Schneider, D. Waithe, S. Galiani, J. Bernardino de la Serna, E. Sezgin, and C. Eggeling, “Nanoscale Spatiotemporal Diffusion Modes Measured by Simultaneous Confocal and Stimulated Emission Depletion Nanoscopy Imaging,” Nano Lett., vol. 18, no. 7, pp. 4233–4240, Jul. 2018, doi: 10.1021/acs.nanolett.8b01190.

[11] G. Vicidomini et al., “STED-FLCS: An Advanced Tool to Reveal Spatiotemporal Heterogeneity of Molecular Membrane Dynamics,” Nano Lett., vol. 15, no. 9, pp. 5912–5918, Sep. 2015, doi: 10.1021/acs.nanolett.5b02001.

[12] V. Mueller et al., “STED nanoscopy reveals molecular details of cholesterol- and cytoskeleton-modulated lipid interactions in living cells.,” Biophys. J., vol. 101, no. 7, pp. 1651–60, Oct. 2011, doi: 10.1016/j.bpj.2011.09.006.

[13] H.-T. He and D. Marguet, “Detecting Nanodomains in Living Cell Membrane by Fluorescence Correlation Spectroscopy,” Annu. Rev. Phys. Chem., vol. 62, no. 1, pp. 417–436, May 2011, doi: 10.1146/annurev-physchem-032210-103402.

[14] C. Eggeling, “STED-FCS Nanoscopy of Membrane Dynamics,” no. August 2012, pp. 291–309, 2012, doi: 10.1007/4243_2012_50.

[15] E. Sezgin, “Super-resolution optical microscopy for studying membrane structure and dynamics,” J. Phys. Condens. Matter, vol. 29, no. 27, 2017, doi: 10.1088/1361-648X/aa7185.

[16] A. Gupta, I. Y. Phang, and T. Wohland, “To Hop or not to Hop: Exceptions in the FCS Diffusion Law,” Biophys. J., vol. 118, no. 10, pp. 2434–2447, 2020, doi: 10.1016/j.bpj.2020.04.004.

[17] S. Veerapathiran and T. Wohland, “The imaging FCS diffusion law in the presence of multiple diffusive modes,” Methods, vol. 140–141, no. 2018, pp. 140–150, 2018, doi: 10.1016/j.ymeth.2017.11.016.

[18] F. Schneider et al., “Diffusion of lipids and GPI-anchored proteins in actin-free plasma membrane vesicles measured by STED-FCS,” Mol. Biol. Cell, vol. 28, no. 11, pp. 1507–1518, 2017, doi: 10.1091/mbc.E16-07-0536.

[19] R. Šachl, J. Bergstrand, J. Widengren, and M. Hof, “Fluorescence correlation spectroscopy diffusion laws in the presence of moving nanodomains (2016 J. Phys. D: Appl. Phys. 49 114002),” J. Phys. D. Appl. Phys., vol. 49, no. 18, p. 189601, May 2016, doi: 10.1088/0022-3727/49/18/189601.

[20] A. Koukalová et al., “Lipid Driven Nanodomains in Giant Lipid Vesicles are Fluid and Disordered,” Sci. Rep., vol. 7, no. 1, p. 5460, Dec. 2017, doi: 10.1038/s41598-017-05539-y.

[21] P. G. Saffman and M. Delbrück, “Brownian motion in biological membranes.,” Proc. Natl. Acad. Sci., vol. 72, no. 8, pp. 3111–3113, Aug. 1975, doi: 10.1073/pnas.72.8.3111.

[22] F. Schneider and E. Sezgin, “Diffusion Measurements at the Nanoscale with STED-FCS,” in Fluorescence Spectroscopy and Microscopy in Biology, M. Šachl, R., Amaro, Ed., Springer, 2022, pp. 323–336. doi: 10.1007/4243_2022_27.

[23] M. Amaro, R. Šachl, G. Aydogan, I. I. Mikhalyov, R. Vácha, and M. Hof, “GM1Ganglioside Inhibits β-Amyloid Oligomerization Induced by Sphingomyelin,” Angew. Chemie - Int. Ed., vol. 55, no. 32, pp. 9411–9415, 2016, doi: 10.1002/anie.201603178.

[24] M. J. Sarmento et al., “The impact of the glycan headgroup on the nanoscopic segregation of gangliosides,” Biophys. J., pp. 1–23, Nov. 2021, doi: 10.1016/j.bpj.2021.11.017.

[25] D. Davidović et al., “Which Moiety Drives Gangliosides to Form Nanodomains?,” J. Phys. Chem. Lett., vol. 14, no. 25, pp. 5791–5797, 2023, doi: 10.1021/acs.jpclett.3c00761.

[26] M. J. Sarmento, J. C. Ricardo, M. Amaro, and R. Šachl, “Organization of gangliosides into membrane nanodomains,” FEBS Lett., vol. 594, no. 22, pp. 3668–3697, Nov. 2020, doi: 10.1002/1873-3468.13871.

[27] E. Sezgin, I. Levental, S. Mayor, and C. Eggeling, “The mystery of membrane organization: Composition, regulation and roles of lipid rafts,” Nat. Rev. Mol. Cell Biol., vol. 18, no. 6, pp. 361–374, 2017, doi: 10.1038/nrm.2017.16.

[28] R. L. Schnaar, R. Gerardy-Schahn, and H. Hildebrandt, “Sialic acids in the brain: Gangliosides and polysialic acid in nervous system development, stability, disease, and regeneration,” Physiol. Rev., vol. 94, no. 2, pp. 461–518, 2014, doi: 10.1152/physrev.00033.2013.

[29] E. Posse de Chaves and S. Sipione, “Sphingolipids and gangliosides of the nervous system in membrane function and dysfunction,” FEBS Lett., vol. 584, no. 9, pp. 1748–1759, 2010, doi: 10.1016/j.febslet.2009.12.010.

[30] L. Canté, M. Corti, D. Acquotti, and S. Sonnino, “Aggregation properties of gangliosides : influence of the primary and secondary structure of the headgroup,” Le J. Phys. IV, vol. 03, no. C1, pp. C1-57-C1-64, May 1993, doi: 10.1051/jp4:1993106.

[31] J. Shi, T. Yang, S. Kataoka, Y. Zhang, A. J. Diaz, and P. S. Cremer, “GM1 clustering inhibits cholera toxin binding in supported phospholipid membranes,” J. Am. Chem. Soc., vol. 129, no. 18, pp. 5954– 5961, May 2007, doi: 10.1021/ja069375w.

[32] S. Arumugam et al., “Ceramide structure dictates glycosphingolipid nanodomain assembly and function,” Nat. Commun., vol. 12, no. 1, pp. 1–12, 2021, doi: 10.1038/s41467-021-23961-9.

[33] I.S. Vinklárek et al., “Experimental Evidence of the Existence of Interleaflet Coupled Nanodomains: An MC-FRET Study,” J. Phys. Chem. Lett., vol. 10, no. 9, 2019, doi: 10.1021/acs.jpclett.9b00390.

[34] R. Šachl, L. B.. Johansson, and M. Hof, “Förster resonance energy transfer (FRET) between heterogeneously distributed probes: Application to lipid nanodomains and pores,” Int. J. Mol. Sci., vol. 13, no. 12, 2012, doi: 10.3390/ijms131216141.

[35] B. Chmelová, J. Humpolíčková, K. Stříšovský, and R. Šachl, “The Analysis of In-Membrane Nanoscopic Aggregation of Lipids and Proteins by MC-FRET,” in Fluorescence Microscopy and Spectroscopy in Biology, R. Šachl and M. Amaro, Eds., 2nd ed.Springer, in press, 2022, pp. 375–400. doi: 10.1007/4243_2022_29.

[36] R. Šachl, J. Humpolíčková, M. Štefl, L. B. Johansson, and M. Hof, “Limitations of electronic energy transfer in the determination of lipid nanodomain sizes,” Biophys. J., vol. 101, no. 11, 2011, doi: 10.1016/j.bpj.2011.11.001.

[37] K. M. Spillane et al., “High-Speed Single-Particle Tracking of GM1 in Model Membranes Reveals Anomalous Diffusion due to Interleaflet Coupling and Molecular Pinning,” Nano Lett., vol. 14, no. 9, pp. 5390–5397, Sep. 2014, doi: 10.1021/nl502536u.

[38] M. Cebecauer et al., “Membrane Lipid Nanodomains,” Chem. Rev., vol. 118, no. 23, pp. 11259–11297, 2018, doi: 10.1021/acs.chemrev.8b00322.

[39] S. Sonnino, E. Chiricozzi, S. Grassi, L. Mauri, S. Prioni, and A. Prinetti, “Gangliosides in Membrane Organization,” 2018, pp. 83–120. doi: 10.1016/bs.pmbts.2017.12.007.

[40] T. Fujiwara, K. Ritchie, H. Murakoshi, K. Jacobson, and A. Kusumi, “Phospholipids undergo hop diffusion in compartmentalized cell membrane.,” J. Cell Biol., vol. 157, no. 6, pp. 1071–81, Jun. 2002, doi: 10.1083/jcb.200202050.

[41] D. M. Andrade et al., “Cortical actin networks induce spatio-temporal confinement of phospholipids in the plasma membrane – a minimally invasive investigation by STED-FCS,” Sci. Rep., vol. 5, no. 1, p. 11454, Jun. 2015, doi: 10.1038/srep11454.

