## Supplementary information for "Revised diffusion law permits quantitative nanoscale characterization of membrane organization"

Barbora Svobodová[a,b], David Šťastný[a,c], Hans Blom[d,e], Ilya Mikhalyov[f], Natalia Gretskaya[f], Alena Balleková[a], Erdinc Sezgin[d,e,*] , Martin Hof[a,*] and Radek Šachl[a,*]

[a] J. Heyrovský Institute of Physical Chemistry of the Czech Academy of Sciences, Dolejškova 3, 182 23 Prague, Czech Republic.

[b] Faculty of Mathematics and Physics, Charles University, Ke Karlovu 5, 121 16 Prague, Czech Republic

[c] Department of Physical and Macromolecular Chemistry, Faculty of Science, Charles University, Hlavova 8, 128 40 Prague, Czech Republic

[d] Science for Life Laboratory, Department of Applied Physics, Royal Institute of Technology, 17165, Solna, Sweden.

[e] Science for Life Laboratory, Department of Women's and Children's Health, Karolinska Institutet, Tomtebodavägen 23, 17165 Solna, Sweden

[f] Shemyakin-Ovchinnikov Institute of Bioorganic Chemistry of the Rusian Academy of Science, 117997, Moscow, Russia

[*]**Correspondence:**

Radek Šachl

Martin Hof

Erdinc Sezgin

### Materials and Methods

#### In-silico generation of STED-FCS diffusion law plots for mobile nanodomains

To generate STED-FCS diffusion law plots for lipid membranes with mobile nanodomains we employed MC simulations. These simulations modelled the two-dimensional movement of lipid molecules within the membranes, generating fluorescence intensity traces for different focal spot sizes, ranging from 10 to 250 nm. The details of this computational approach are described in [1][2]. This approach mimics the outcomes of a typical STED-FCS experiment. Subsequently, we auto-correlated these fluorescence traces in the same manner as experimentally acquired data to derive STED-FCS autocorrelation functions for different focal spot sizes ($G(\tau, w)$):

$$G(\tau, w) = \frac{\langle \Delta I(t) \Delta I(t+\tau) \rangle}{\langle \Delta I(t) \rangle^2}. \tag{1}$$

Here $I$(t) represents intensity at time $t$ and $\tau$ is the lag-time. To determine the probe diffusion coefficient $D(w)$ for each spot size, we fitted these autocorrelation curves using a model that considers anomalous two-dimensional diffusion:

$$G(\tau) = 1 + \frac{1}{N} \cdot \frac{1}{(1 + \frac{\tau}{\tau_D})^{\alpha}} \tag{2}$$

Here, $\tau_D$ is the diffusion time, representing the average time for a molecule to move from the center to the edge of the focal spot, and it is related to $D$ via $D = \frac{w^2}{4\tau_D}$. In equation 2, $N$ denotes the number of particles within the analyzed volume, and $\alpha$ is the anomalous factor. For free diffusion, $\alpha$ = 1, but as diffusion deviates from this ideal, $\alpha$ deviates accordingly [3], [4]. Finally, we constructed a set of STED-FCS diffusion law plots by plotting $D$ versus $w$, combining the parameters outlined in **Table** Chyba! Nenalezen zdroj odkazů.**S1** to depict the dependence of the diffusion law plot shape on these variables.

**Table S1:** Input parameters used for Monte Carlo simulations of probe diffusion in the presence of moving nanodomains. The parameters involve probe diffusion coefficient outside $D_{out}$ and inside $D_{in}$ nanodomains, nanodomain radius $R_d$, membrane surface fraction occupied by nanodomains and the probe distribution coefficient between nanodomains and the surroundings $K_d$. The scheme illustrates the process of parameter variation, with a separate simulation conducted for each listed combination in the table. Parameters highlighted by red squares remained unchanged throughout the scheme unless their specific impact was under investigation. The values for nanodomain diffusion $D_d$ were calculated according to the Saffman- Delbrück model [5].

| Par | Value | Diagram |
|---|---|---|
| $D_{out}$ ($\mu m^2/s$) | 9 | |
| $R_d$ (nm) | 25 – 120 | |
| $D_d$ ($\mu m^2/s$) | 2.6 – 4.4 | |
| $D_{in}$ ($\mu m^2/s$) | 0.9 – 9 | |
| $K_d$ (-) | 2.5 – 50 | |
| $f$ (-) | 10 – 50 | |

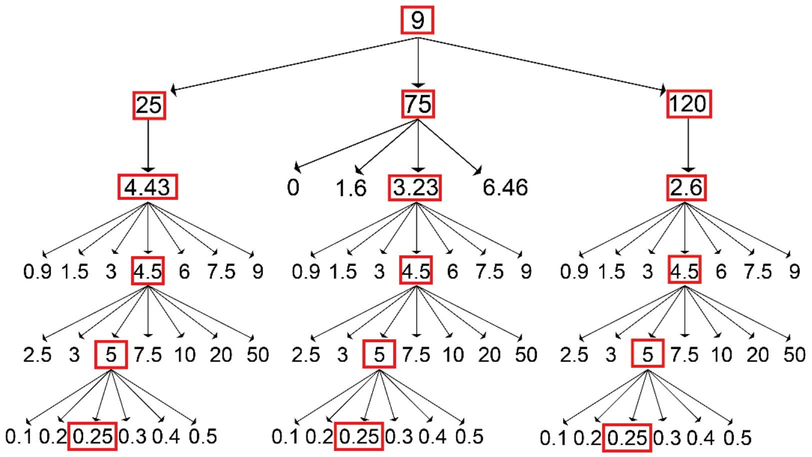

#### Quantitative analysis of STED-FCS diffusion law plots

For the quantitative analysis of the STED-FCS diffusion law plot dependencies, we generated a comprehensive set of diffusion law plots for different combinations of simulation input parameters, including $D_d$, $D_{in}$, $K_d$, $R_d$ and $f$. The complete library of these generated dependencies is available at the following link: **[data repository link forthcoming]** and can be used to fit experimentally obtained STED-FCS diffusion law plots. In these simulations, we specifically considered the diffusion of lipid probes outside of the nanodomains $D_{out} = 9$ $\mu m^2/s$, corresponding to free probe diffusion in a homogeneous DOPC/Chol membrane. In cases where $D_{out}$ takes other values, all diffusion coefficients can be normalized so that the values of all diffusion coefficients are relatively conserved.

Our fitting approach involved comparing experimentally measured diffusion law plots with the generated diffusion dependencies by calculating a chi-squared parameter to characterize their similarity. We analysed the chi-squared values only for the physically realistic combinations of all parameters. For the diffusion dependencies of GUVs, where $R_d$ and $f$ were known a priori from independent MC-FRET analysis, we explored only the combinations of $D_d$, $D_{in}$, and $K_d$. In the final stage, slight adjustments were made to the best-matching curve to optimize its vertical positioning, involving fine-tuning the fraction $f$.

In the case of GPMVs and cell plasma membranes, the situation differed because neither the size of the nanodomains nor the fraction of surface coverage by nanodomains was known in advance. Therefore, in this case, we primarily focused on the determination of these two characteristic parameters. Based on the GUV experiments (see the Result section), we made the following assumptions: 1) the nanodomains move according to the Safmann-Delbrück model, where $D_d$ depends on the size of the nanodomains and the mobility of the molecules in the surrounding environment ($D_{out}$). 2) $GM_1$-Atto565 has a relatively high affinity for existing nanodomains, characterized by $K_d = 5$. Finally, 3) the mobility of $GM_1$-Atto565 within nanodomains is half that of the mobility in the surrounding non-domain phase. These assumptions allowed us to reduce the number of optimized parameters to $R_d$ and $f$, leaving $D_d$, $K_d$ and $D_{in}$ fixed. We also attempted to optimize $D_{in}$ in the final step, but this optimization did not improve the fit in the investigated range of physically acceptable $D_{in}$ values (from $D_{in} = D_{out}/3$ to $D_{in} = D_{out}/1.5$), confirming $D_{in} = D_{out}/2$.

#### STED-FCS experiments on GUVs: methodology

STED-FCS data were taken at a Leica FALCON microscope. The $GM_1$-Atto565 probes were excited using a 561 nm white light laser and the fluorescence was depleted using a tuneable red laser at 660 nm. The emission was recorded at 575-635 nm. Fitting was performed with FoCuS-point fitting program[6]. For calibration of the spot size, measurements of DOPC GUVs with different STED laser powers were utilized. Assuming free diffusion allows the spot size calculation according to the following equation:

$$w_{\mathrm{STED}} = w_{\mathrm{confocal}}\sqrt{\tau_{\mathrm{STED}}/\tau_{\mathrm{confocal}}}\ , w_{conf} = \sqrt{D4\tau_{\mathrm{confocal}}}$$

where $w$ is the radius of the spot size and $\tau_{\mathrm{confocal}}$, $\tau_{\mathrm{STED}}$ the transit times which the molecule spends in the observation spot in confocal and at a certain STED laser power, respectively. Individual ACFs were fitted with eq.2.

#### Materials

1,2-dioleoyl-sn-glycero-3-phosphocholine (DOPC), $GM_1$ ganglioside (bovine brain sodium salt), Nstearoyl-D-erythro-sphingosylphosphorylcholine (SM) and cholesterol (ovine wool) were purchased from Avanti Polar Lipids (Alabaster, AL, USA). Sucrose, Bovine serum albumin (BSA) and Phophate Buffered saline (PBS) were purchased from Sigma Aldrich (St.Louis, USA). Organic solvents of spectroscopic grade were purchased from Merck (Darmstadt, Germany). Fluorescent probe Atto565 was coupled to $GM_1$ as described below.

#### Preparation of GUVs

Giant unilamellar vesicles (GUVs) were prepared using electroformation in custom-made Teflon chambers equipped with two platinum electrodes [7]. Lipids were dissolved in either chloroform or a 2:1 chloroform/methanol mixture, with fluorescently labeled ganglioside GM1 (GM1-Atto565) added at a probe-to-lipid ratio of 1:5000. A 6 µl volume of this lipid mixture was spread on each electrode and allowed to evaporate. Then, 370 µl of a sucrose solution (300 mOsm/kg) was added to the chamber. The electroformation protocol consisted of applying an alternating electric field (2 V peak-to-peak, 10 Hz) for 1 hour while incubating the samples at 47°C. Afterward, the frequency was reduced to 2 Hz, and the electric field (2 V peak-to-peak) was applied for an additional 30 minutes. The samples were then allowed to cool gradually. In the final steps, 150 µl of BSA was used to coat an 8-well µ-Slide (Ibidi, Munich, Germany). Following this, 50 µl of GUVs and 100 µl of PBS buffer were added to the slide.

#### Tagging Atto565 to $GM_1$

$GM_1$-Atto565 was obtained by re-acylation of de-acetylated $GM_1$, received as described in [8]. In brief, 4 mg (2.6 nmol) de-acetylated $GM_1$ was dissolved in 1 ml of freshly distilled DMF, 1 ml of triethylamine was added and 2 mg (2.9 nmol) of Atto565-N-hydroxysuccinimidyl ester (Sigma-Aldrich), dissolved in 200 µl of MeCN was added. The reaction mixture was stirred on a magnetic stirrer for 96 h at room temperature. The reaction mixture was evaporated and $GM_1$-Atto565 was isolated by column chromatography, using Silica gel 100 (Merck) in chloroform-methanol-water, 65:25:4 (v/v/v) with a yield of 1.1 mg (0.52 nmol) (14%, $M_r$ = 1997). $R_f$ = 0.45 (TLC chloroform : methanol : $H_2O$, 60:40:9, v/v/v), where $R_f$ is substance's mobility in the TLC chromatography, calculated as the fraction of the front mobility.

### Supporting results:

| Ratiometric parameter | | 160/60 | | | $R_{confin}$ | | | $S$ | | |
|---|---|---|---|---|---|---|---|---|---|---|
| Focal spot size (nm) | | 60-160 | | | 60 | | | 60-160 | | |
| Domain size | | Large | Intermediate | Small | Large | Intermediate | Small | Large | Intermediate | Small |
| Changing $D_{in}$ | 0.9 | 2.02 | 1.73 | 1.15 | 5.46 | 2.82 | 1.71 | 16.78 | 23.20 | 7.73 |
| | 1.5 | 1.95 | 1.76 | 1.08 | 5.17 | 2.74 | 1.59 | 16.45 | 25.10 | 4.63 |
| | 3 | 1.74 | 1.49 | 1.12 | 4.51 | 2.14 | 1.42 | 14.69 | 20.79 | 7.69 |
| | 4.5 | 1.66 | 1.45 | 1.09 | 3.78 | 1.81 | 1.27 | 15.80 | 22.32 | 6.53 |
| | 6 | 1.65 | 1.43 | 1.08 | 3.63 | 2.01 | 1.32 | 16.06 | 19.03 | 5.76 |
| | 7.5 | 1.33 | 1.29 | 1.08 | 2.75 | 1.72 | 1.26 | 10.86 | 15.28 | 5.73 |
| | 9 | 1.21 | 1.27 | 1.03 | 2.34 | 1.57 | 1.18 | 8.02 | 15.45 | 2.46 |
| Changing $K_d$ | 2.5 | 1.19 | 1.20 | 1.07 | 1.62 | 1.26 | 1.12 | 10.41 | 14.31 | 5.51 |
| | 3 | 1.22 | 1.32 | 1.06 | 2.14 | 1.58 | 1.22 | 9.07 | 18.06 | 4.32 |
| | 5 | 1.55 | 1.40 | 1.09 | 3.24 | 2.02 | 1.31 | 15.27 | 17.68 | 6.13 |
| | 7.5 | 2.00 | 1.65 | 1.02 | 4.50 | 2.26 | 1.37 | 20.01 | 25.87 | 1.20 |
| | 10 | 2.35 | 1.71 | 1.10 | 5.43 | 2.58 | 1.53 | 22.37 | 24.62 | 5.62 |
| | 20 | 2.56 | 1.75 | 1.04 | 6.72 | 2.95 | 1.66 | 20.89 | 22.89 | 2.42 |
| | 50 | 2.70 | 1.79 | 1.01 | 7.70 | 3.15 | 1.81 | 19.91 | 22.74 | 0.58 |
| Changing $f$ | 10% | 1.29 | 1.41 | 1.15 | 3.91 | 2.48 | 1.35 | 5.94 | 14.76 | 10.10 |
| | 20% | 1.46 | 1.41 | 1.14 | 3.57 | 2.15 | 1.34 | 11.54 | 17.17 | 9.37 |
| | 30% | 1.64 | 1.48 | 1.03 | 3.29 | 1.85 | 1.37 | 17.56 | 23.35 | 2.21 |
| | 40% | 1.79 | 1.38 | 0.96 | 2.66 | 1.69 | 1.48 | 26.87 | 20.30 | -2.13 |
| | 50% | 1.67 | 1.26 | 0.94 | 2.18 | 1.62 | 1.70 | 27.56 | 14.21 | -3.23 |

**Figure S1:** An extended version of **Figure 3** presenting ratiometric parameters $D_{\mathrm{confocal}}/D_{\mathrm{STED}}$ represented by the red color map, $R_{\mathrm{confin}}$ by the violet colour map, and $S$ by the yellow colour map used as indicators of nanodomain size and fraction. The parameters were calculated for large ($R_d$ = 120 nm), intermediate ($R_d$ = 75 nm), and small nanodomains ($R_d$ = 25 nm), based on the probe diffusion coefficient within the nanodomains ($D_{in}$), probe partition coefficient between nanodomains and surroundings ($K_d$), and fraction of nanodomains in the membrane ($f$). If not stated otherwise, $D_{in}$ = 4.5µm$^2$/s, $K_d$ =5, $f$ = 0.25. The colour code illustrates a gradual change from low (light) to high (dark) values.

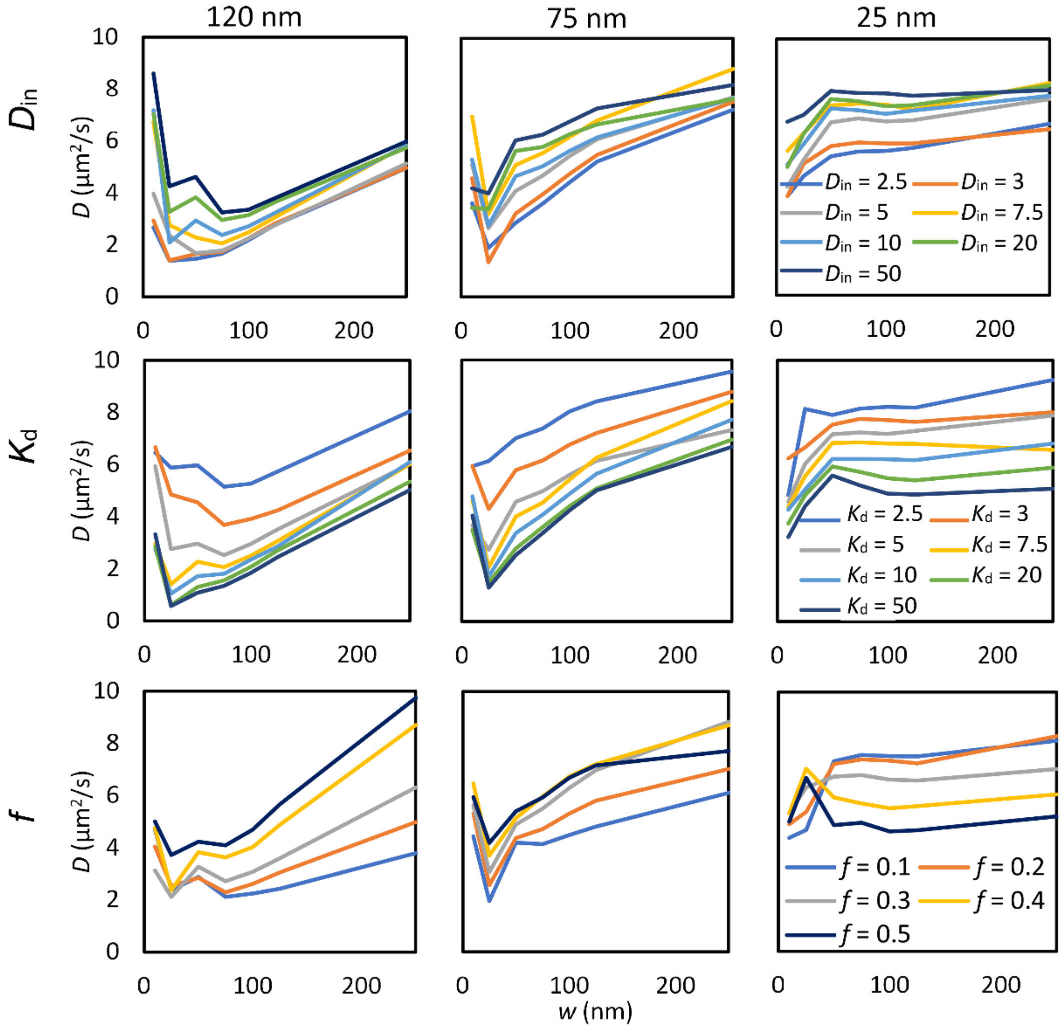

**Figure S2:** Computationally generated STED-FCS diffusion law plots illustrating mobile nanodomains with $D_d$ modelled according to the Saffman-Delbrück model. Plots are shown for three nanodomain sizes: large (left column), intermediate (middle column) and small (right column). The impact of the probe diffusion coefficient within the nanodomains ($D_{in}$) is depicted in the upper row, the probe distribution coefficient between the nanodomains and the surroundings ($K_d$) in the middle row, and the area fraction ($f$) occupied by nanodomains in the lower row. If not stated otherwise, $D_{in}$ = 4.5µm²/s, $K_d$ =5, $f$ = 0.25.

#### Trends in the anomalous coefficient alpha

Building upon the analysis presented in **Figure S2**, we extended our investigation to establish a relationship between the anomaly factor (referred to as alpha) and the size of the confocal spot $w$, termed alpha plot dependency (**Figure S3**). This factor, determined by fitting individual STED-FCS autocorrelation functions by a model accounting for anomalous diffusion (**Eq 2**), equals 1 for free diffusion, while it achieves values less than 1 for anomalous diffusion. Importantly, the resulting dependencies closely resemble the diffusion law plots outlined in **Figure S2**, exhibiting a distinct asymmetric funnel shape. This characteristic shape thus emerges as an additional indicator of nanodomain size. A potential drawback of this approach might be that achieving the minimum alpha necessitates extremely small confocal spot sizes, which may pose challenges in STED microscopy.

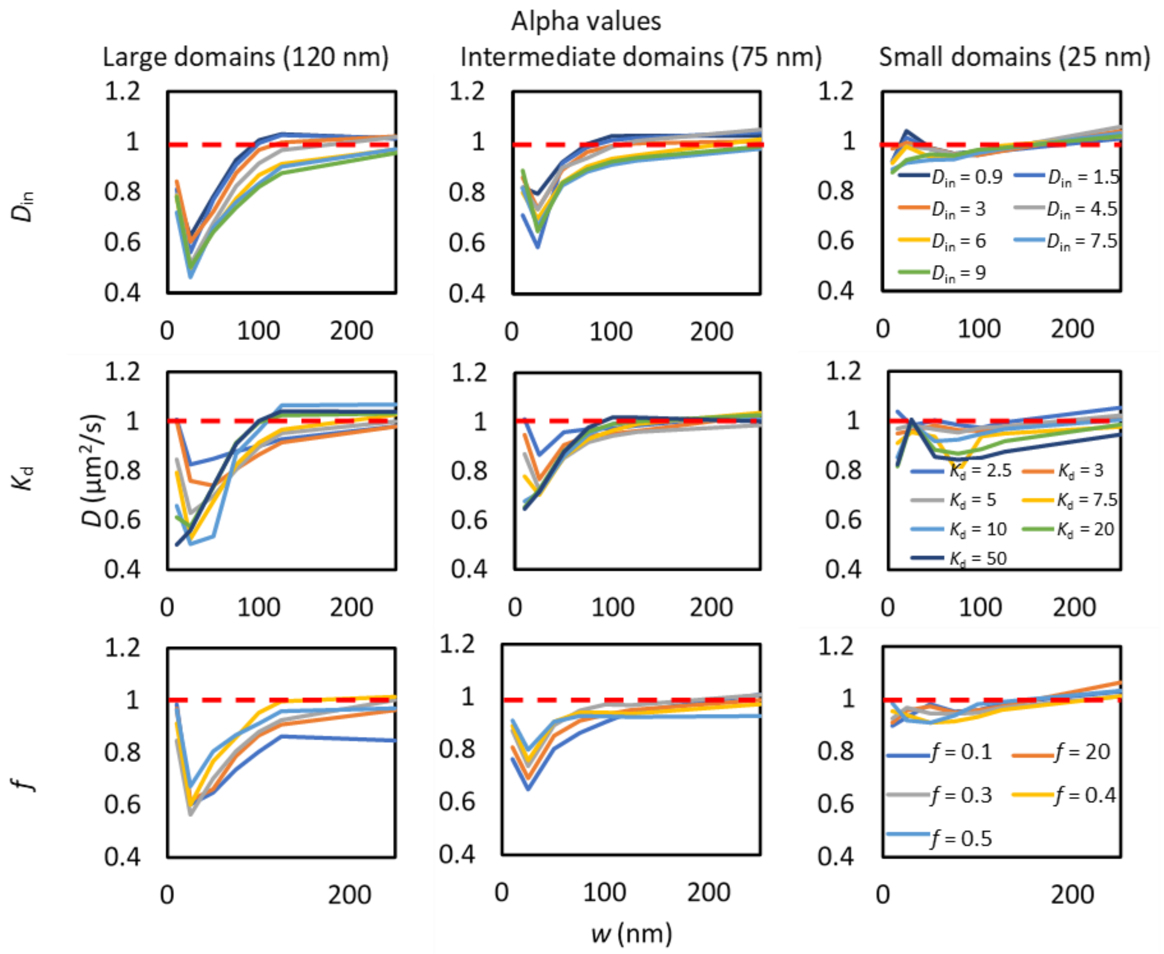

**Figure S3:** A set of alpha plots complementary to **Figure S2** establishing a relationship between the anomaly factor α and the size of the confocal spot *w*. Alpha plot dependency exhibits a pronounced funnel shape, whereby the depth of this funnel's minimum increases with larger moving nanodomains. For nanodomains with $R_d$ = 120 nm, the minimum alpha falls within the range of 0.4-0.6, while for nanodomains with $R_d$ = 75 nm, the minimum alpha rises to 0.6-0.8. Nanodomains with $R_d$ = 25 nm show a slight decrease in alpha, ranging from 0.9 to 1. The impact of Din is depicted in the upper row *K*d in the middle row, and f lower row. If not stated otherwise, $D_{in}$ = 4.5 µm$^2$/s, $K_d$ =5, $f$ = 0.25.

**Reference:**

[1] L. Wawrezinieck, H. Rigneault, D. Marguet, and P. F. Lenne, "Fluorescence correlation spectroscopy diffusion laws to probe the submicron cell membrane organization," *Biophys. J.*, vol. 89, no. 6, pp. 4029–4042, 2005, doi: 10.1529/biophysj.105.067959.

[2] R. Šachl, J. Bergstrand, J. Widengren, and M. Hof, "Fluorescence correlation spectroscopy diffusion laws in the presence of moving nanodomains (2016 J. Phys. D: Appl. Phys . 49 114002)," *J. Phys. D. Appl. Phys.*, vol. 49, no. 18, p. 189601, May 2016, doi: 10.1088/0022-3727/49/18/189601.

[3] E. Sezgin *et al.*, "Partitioning, diffusion, and ligand binding of raft lipid analogs in model and cellular plasma membranes," *Biochim. Biophys. Acta - Biomembr.*, vol. 1818, no. 7, pp. 1777–1784, 2012, doi: 10.1016/j.bbamem.2012.03.007.

[4] E. Sezgin *et al.*, *Measuring nanoscale diffusion dynamics in cellular membranes with super-resolution STED–FCS*, vol. 14, no. 4. 2019.

[5] P. G. Saffman and M. Delbrück, "Brownian motion in biological membranes.," *Proc. Natl. Acad. Sci.*, vol. 72, no. 8, pp. 3111–3113, Aug. 1975, doi: 10.1073/pnas.72.8.3111.

[6] D. Waithe, M. P. Clausen, E. Sezgin, and C. Eggeling, "FoCuS-point: Software for STED fluorescence correlation and time-gated single photon counting," *Bioinformatics*, vol. 32, no. 6, pp. 958–960, 2016, doi: 10.1093/bioinformatics/btv687.

[7] M. I. Angelova and D. S. Dimitrov, "Liposome electroformation," *Faraday Discuss. Chem. Soc.*, vol. 81, pp. 303–311, 1986, doi: 10.1039/DC9868100303.

[8] S. Sonnino, G. Kirschner, R. Ghidoni, D. Acquotti, and G. Tettamanti, "Preparation of GM1 ganglioside molecular species having homogeneous fatty acid and long chain base moieties.," *J. Lipid Res.*, vol. 26, no. 2, pp.

248–257, Feb. 1985, doi: 10.1016/S0022-2275(20)34395-9.
